# Scaffold-Free Acoustic Levitation Platforms Enable Scalable Culture of Neuronal Spheroids and Assembly of Layered Cortico - Striatal Assembloids

**DOI:** 10.64898/2026.04.02.716069

**Authors:** Chloé Dupuis, Guillaume Viraye, Xavier Mousset, Nathan Jeger-Madiot, Jean-Luc Aider, Jean-Michel Peyrin

## Abstract

Engineering three-dimensional neuronal tissues with defined architecture and functional connectivity remains a critical challenge for applications in disease modeling, drug discovery, and regenerative medicine. Recently, a variety of fabrication methods have arisen, such as bioprinting or manual assembly of organoids, but often struggle with scalability, reproducibility, or maintaining cell viability. Here, two scaffold-free acoustic levitation bioreactors are introduced: one optimized for the culture of uniform neuronal spheroids, and another designed for the structuration of assembloids composed of distinct neuronal identities. Using acoustic standing waves, these platforms enable the contactless manipulation of cells and aggregates, facilitating the formation of highly viable functionally mature spheroids. This study shows that both striatal and cortical cell aggregates formed in acoustic levitation self-organize into spheroids within 24 hours and remain viable up to 10 days under these particular culture conditions without medium renewal. These neuro-spheroids demonstrate healthy development with increased growth and typical terminal differentiation and synaptic maturation. Moreover, concentric cortico-striatal assembloids were successfully structured and cultivated using optimized acoustofluidic chips. Offering versatile and scalable tools for engineering complex neuronal networks, acoustic levitation reveals itself as an innovative approach to 3D neuronal tissue modeling, with broad implications for bioengineering, regenerative medicine and fundamental neuroscience research.

## 1 Introduction

The development of three-dimensional (3D) microphysiological systems has revolutionized our ability to model the nervous system, offering unprecedented opportunities to study brain development, disease mechanisms, and therapeutic interventions. Traditional approaches to engineering neuronal tissues have relied on either top-down or bottom-up strategies. Top-down methods aim to recapitulate the brain’s developmental processes by guiding stem cells through differentiation pathways using spatially and temporally controlled biochemical and biophysical cues, such as growth factor gradients or mechanical constraints. These approaches enable the formation of complex, region-specific tissue architectures that closely mimic in vivo morphogenesis [1, 2]. In contrast, bottom-up strategies assemble neuronal networks from modular components —such as cells, spheroids, organoids, or microtissues— using advanced bioengineering techniques. This modularity allows for precise control over tissue composition, structural organization, and connectivity, making it particularly suitable for scalable and customizable applications in disease modeling and drug screening [3].

Despite these advances, significant challenges remain in constructing neuronal tissues with defined topologies and functional connectivity. Bioprinting, for example, enables the precise spatial arrangement of cells and biomaterials, yet struggles with maintaining high cell viability and achieving the resolution necessary for complex neural architectures [4, 5]. Another promising approach involves the assembly of assembloids —complex structures formed by combining distinct organoids or spheroids— using chemical cues or manual techniques. While assembloids can model interactions between different brain regions, their formation often lacks reproducibility and scalability, limiting their utility for studying long-distance neuroanatomical pathways [6, 7, 8].

To address these limitations, we introduce a scaffold-free, acoustic levitation-based platform for the structured culture and assembly of neuronal assembloids. Acoustic levitation has emerged as a powerful tool in tissue engineering, enabling the contactless manipulation and patterning of cells and cell aggregates without the need for external scaffolds or hydrogels [9, 10]. Previous studies have demonstrated its potential for levitating and culturing mammalian cells [11, 12, 13], patterning cell aggregates in hydrogels [14, 15, 16], and even promoting the self-organization of stem cells into spheroids [17]. Notably, acoustic levitation has been shown to enhance the functionality of hepatocytes compared to conventional culture methods [18, 19, 20], and to enable the controlled fusion of spheroids and organoids [21, 22, 23].

In this work, we present two distinct acoustic levitation bioreactors: one designed for the culture of multiple neuronal spheroids (Figure 1.a-c), and another for the precise structuration and assembly of assembloids composed of distinct neuronal identities (Figure 6.d-e). While we demonstrate that acoustic levitation allows instant formation and long term culture of viable neuronal spheroids, we also show that acoustic waves transiently increase neuroprecursors proliferation leading to fully differentiated neuro-spheroids. Moreover, using a tuned acoustofluidic approach, we show that 3D layered cortico-striatal neurospheroids can be constructed. This enables reproducible and scalable reconstruction of neuro-anatomical pathways, offering a versatile tool for engineering functional neuronal networks with precise spatial organization. By integrating acoustic levitation with advanced bioengineering principles, we provide a transformative approach for 3D neuronal tissue modeling, with broad implications for healthcare, from fundamental neuroscience research to drug discovery and regenerative medicine.

**Figure 1:**
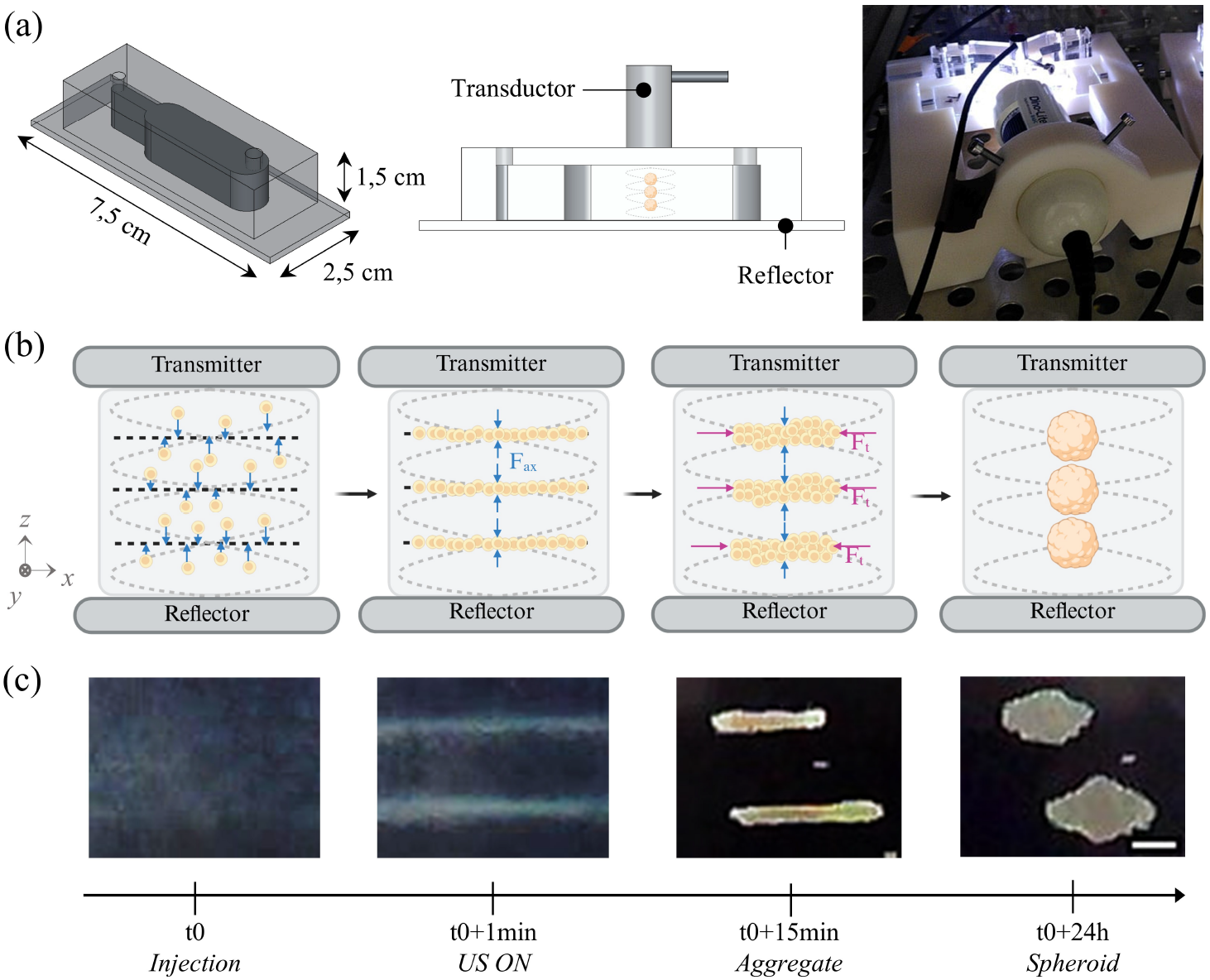
Formation and culture of neurospheres in acoustic levitation. (a) Multi-node bulk acoustic standing wave resonator composed of a PDMS cavity with on one side a transmitter producing ultrasonic waves and on the other side a reflector allowing for the emergence of the stationary field. A picture on the right shows the acoustofluidic chip inside a 3D printed support allowing for precise positioning of the transducer and the camera to monitor formation and self-organization of the spheroids. (b) Schematics describing the aggregation of cells and formation of spheroids in acoustic levitation. When cells are injected in a multi-node cavity, they aggregate at the pressure nodes highlighted by bold black dotted lines. The acoustic pressure is represented by gray dotted curves with the pressure being null at the pressure nodes. The axial component of the ARF is represented with blue arrows and the transversal component is represented with magenta ones. The resonance condition is met as the height of the cavity corresponds to a multiple of half the wavelength of the ultrasonic wave. Here, to create 3 pressure nodes with an ultrasound wave of 2 MHz frequency, corresponding to ≈ 750 *µ*m wavelength in water, the cavity is ≈ 1.2 mm high. Schematics were created with Biorender.com. (c) Timeline with focused snapshots illustrating the formation of two neuronal spheroids inside a multi-node acoustofluidic. After injection and homogenization of the cell suspension inside the cavity, it takes roughly one minute for the ultrasonic stationary field to stabilize and produce the axial ARF that pushes cells towards the pressure nodes. Then, the radial ARF drives them at the center of the cavity where they aggregate in a flat disk shape. During the first 24h, cells start to adhere to each other and reorganize into a spheroid, trapped inside the pressure node. Scale bar is 150 µm.

## 2 Results

### 2.1 Self-organization of cortical and striatal spheroids in acoustic levitation

First, a multi-node acoustofluidic chip was developed to study the self-organization and viability of multiple neuronal spheroids in acoustic levitation. Objects in suspension in a fluid medium can be handled without contact by ultrasound through the Acoustic Radiation Force (ARF), which arises when acoustic waves propagate through a medium and transfer their momentum to the objects. In this study, the ARF was created by an acoustic standing wave in acoustofluidic chips containing a resonant cavity (Figure 1.a). In practice, an acoustic wave is emitted by an ultrasonic transducer (located on top of the chip) and propagates inside the cavity through a transmitting layer made of PDMS (Polydimethylsiloxane). The transmitted wave then propagates through the fluid medium toward the opposite wall, the reflector (on the bottom of the chip), which then reflects an important part of the wave toward the emitter. The resonance condition requires that the cavity height *h* is a multiple of half the acoustic wavelength 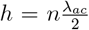. If this condition is satisfied, then all particles or cells suspended in the fluid are moved toward the closest acoustic pressure node by the axial component of the ARF, *F*_*ax*_, as illustrated in Figure 1.b. Once in the acoustic pressure node, also called the acoustic levitation plane, the axial component of the ARF vanishes and the cells are moved radially toward the center of the cavity by the radial component of the ARF *F*_*t*_, which is maximum in the acoustic levitation plane [24, 25]. Because of the short distance interaction between the objects induced by the Bjerknes force, the aggregated particles form a cohesive aggregate which can be maintained in acoustic levitation as long as needed [26, 27, 28, 29].

For cell culture, the fluid medium used was neuronal cell culture medium and the acoustic contrast factor of the cells was then positive [30, 31], leading to the formation of cells layers in the pressure nodes. As illustrated in Figure 1., after a few seconds, cells moved towards the pressure nodes and organized in layers. After 15 min, it leads to the formation of cohesive cell aggregates and after a few hours, primary brain cells self-organized into spheroids as previously observed with other cell types [17, 18]. Neurospheres were then kept in acoustic levitation for several days.

The self-organization of primary cells extracted from two different brain regions was monitored over long-term culture using a compact digital microscope on the side of the chip (Figure 1.a). The morphological evolution of both cortical (CxSs) and striatal (StSs) spheroids was quantified using image analysis and geometric morphodescriptors (circularity and eccentricity) described in the Methods section. Initially, cortical and striatal progenitors were organized in roughly disk-like layers of equivalent diameter *D*_*e*_ ranging from 150 to 250 *µ*m. A 2 MHz ultrasound source was used, which corresponds to an acoustic wavelength *λ*_*ac*_ ≈ 750 *µ*m in water, meaning that the axial separation between two cell layers is 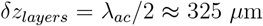. This also defines the size limit of the objects that can be handled in acoustic levitation with this ultrasound frequency. In 8 mm high multi-node acoustofluidic chip, 6 spheroids can be cultivated in acoustic levitation in parallel.

As already observed with other cell types in similar acoustofluidic chips [17, 18, 32], primary neurons started forming layers quickly (a few minutes) before adhering to each other and self-organizing during the first 24 hours. This can be seen with the increase of the equivalent diameter *D*_*e*_ from *t* = 0 to *t* = 10^0^*days* ≡ 24*h* on figure 2.b. As a result of this adhesion mechanism and cell-cell interactions, aggregates changed from a layer-like shape to a more ellipsoidal shape before reaching an equilibrium as spheroids. At the end of these steps of self-organization, spheroids formed in acoustic levitation reached an average equivalent diameter ⟨*D*_*e*_⟩ of 210 *µm* for StSs and of 280 *µm* for CxSs. During the following days of culture in acoustic levitation, the sizes of spheroids remained stable for both subtypes of primary neurons.

**Figure 2:**
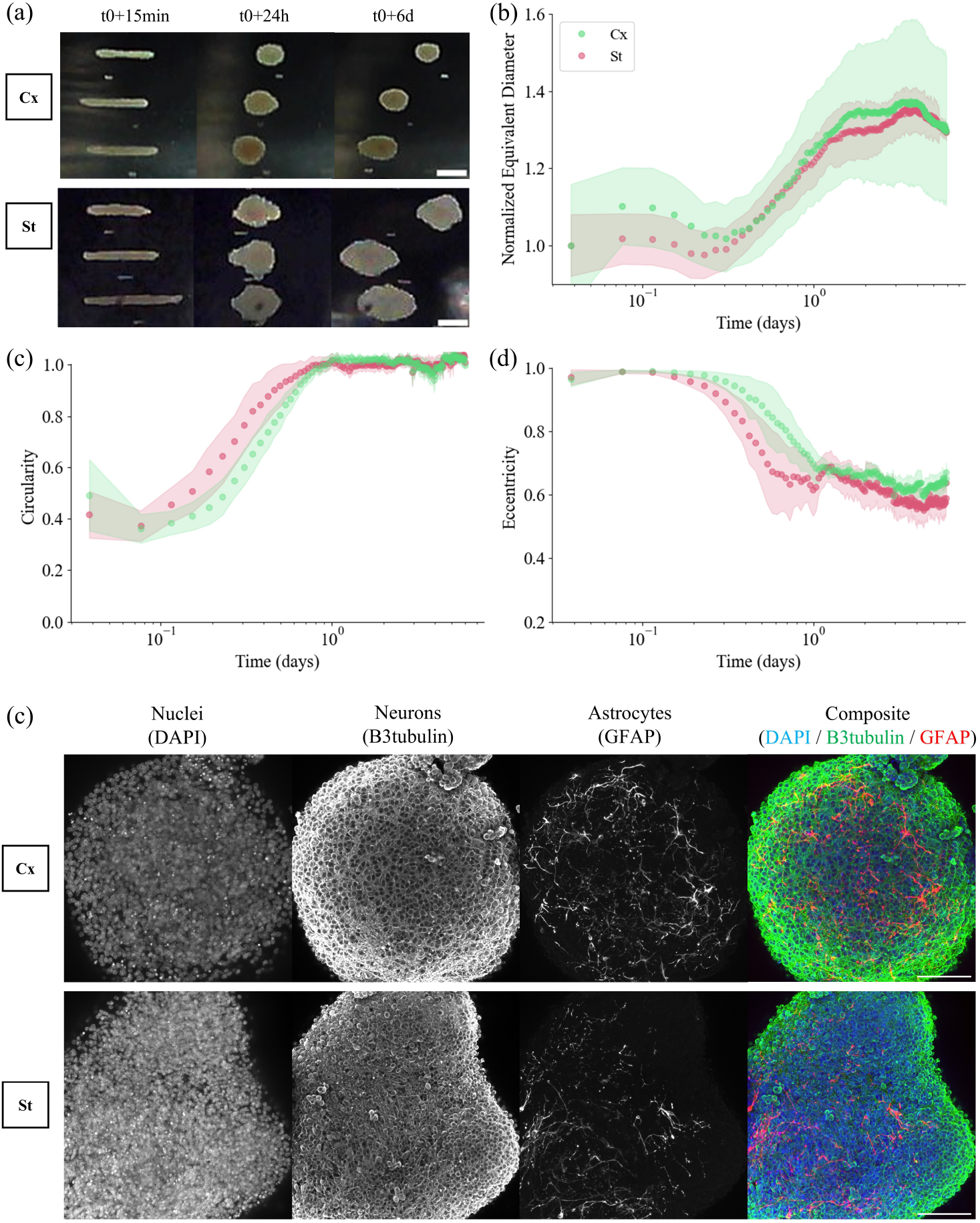
Morphodynamic analysis of spheroids formation. (a) Time-lapse of both cortical (Cx) and striatal (St) primary brain cells aggregates in acoustic levitation, monitored from the side of the *Multi-Node Chip* and cultivated for 6 days. Images were taken every 5 minutes with a *Dinolite* camera, inside a conventional incubator (setup shown in Supplementary Data). Whole time-lapses of both CxSs & StSs are available on the Supplementary Video 1 & 2 resp. Scale bar is 150 µm. (b) Time evolution of the equivalent diameter *D*_*e*_(*t*), computed to evaluate the apparent size of the spheroids. Circularity and eccentricity are shown in (c) and (d) respectively. Average values are represented by dots while shades of color highlight the standard deviation. CxSs are plotted in green (n=6) and StSs in red (n=5). Time is in logarithmic scale to show in more details the onset of self-assembly in the first 24 hours of culture. (e) Projection of immunofluorescence confocal images of 6 days old cortical (CxS) and striatal spheroids (StS) cultivated in acoustic levitation showing nucleus (DAPI), axons of neurons (B3tubulin) and astrocytes intermediate filaments (GFAP). Scale bar is 100 *µ*m.

Figure 2.c shows the time evolution of circularity and eccentricity for cortical and striatal spheroids. They both reorganized into spheroids after the first 24 hours of culture in acoustic levitation and kept this shape afterward. The dynamic of self-organisation seems greater for striatal spheroids, meaning StSs reached the spherical state before CxSs. This phenomenon can also be observed in figure 2.d as the eccentricity also decreases faster for StSs than for CxSs, both reaching a value of 0.67 at the end of the first 24 hours. Another interesting point is that both spheroids kept their circularity close to 1 after 24h, with little variations during the rest of the culture. On the other hand, the eccentricity of StSs shows larger fluctuations than CxSs after 24h, as illustrated by the relatively large standard deviation. In addition, the average eccentricity of StSs continues to decrease even after 24 h, reaching a final value of 0.57 after 6 days of culture. This is also qualitatively illustrated on timelapse snapshots shown in Figure 2.a. The StSs usually presented a *“rougher”* surface, with small buds, whereas CxSs had a relatively smooth circular shape.

After 6 days in acoustic levitation, cortical and striatal spheroids were retrieved and immuno-stained before being imaged with a confocal microscope. As can be seen from the projections of the full-mount confocal images in figure 2.e, neurospheres of the two subtypes maintained high viability and neuroprogenitors differentiated both in mature neurons - with extended axons - and in astrocytes showing a healthy phenotype.

### 2.2 Viability of cortical and striatal spheroids cultivated in acoustic levitation

The viability of the spheroids of both neuronal subtypes cultivated in acoustic levitation was then assessed in more detail. Control spheroids were cultured in commercial multi-well plates and without continuous exposure to acoustic waves. Neurospheres being dense and cohesive, analysis pipelines based on histological studies were developed (see Methods). Shortly, immunofluorescence staining were performed on thin cryosectioned slices of spheroids to access cell proliferation, survival, morphology and phenotype throughout the spheroids. Viability assessment was based on the morphology of DAPI labeling as it was shown to be related with cell viability, since polynucleolated nuclei are associated with healthy cells while small nuclei with intense signal correspond to condensed DNA of apoptotic cells [33].

Both control and levitated spheroids demonstrated good cell survival after 3 days, with a high majority of cells presenting polynucleolated nuclei, homogeneously distributed throughout cortical and striatal spheroids. Overall, only few small clusters or single cells showed diffuse and/or intense labeling, attributed to apoptotic state, for both levitated and control conditions (figure 3.a & c). The percentages of polynucleolated nuclei over the total number of nuclei, i.e. the spheroid survival rates, were calculated for the different conditions on day 3, 6 and 10 and are shown for cortical spheroids on figure 3.b and striatal spheroids on figure 3.e. For both subtypes and conditions, the viability decreases over time, which was expected from natural developmental apoptosis as well as the lack of medium renewal. After 3 days of culture, both levitated and control CxSs and StSs have similar survival rates of almost 90%. With increasing duration of culture, both levitated CxSs and StSs show a higher survival rate than the control ones.

**Figure 3:**
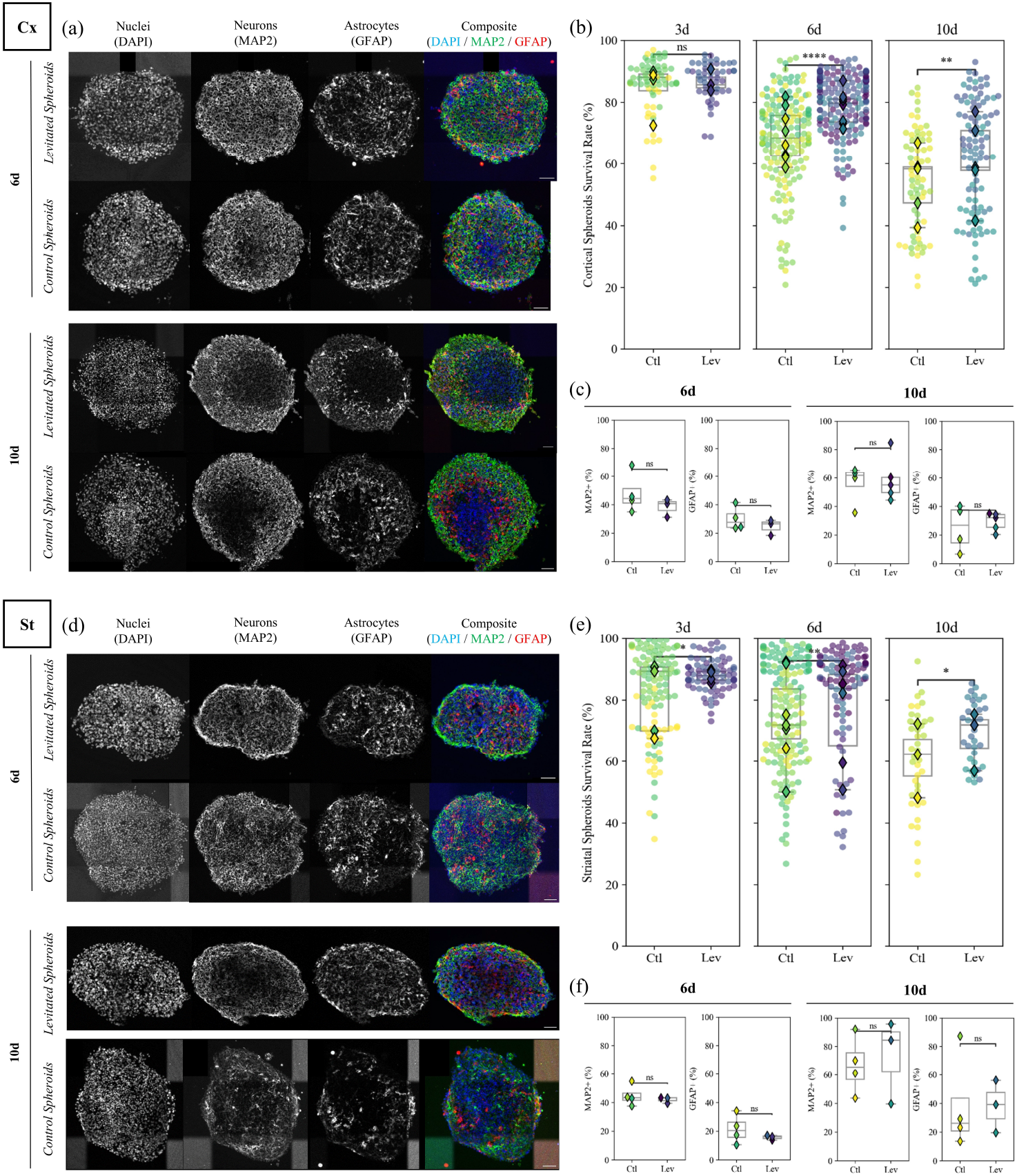
Characterization of survival and phenotypic maturation of cortical (CxSs) and striatal (StSs) spheroids. Immunofluorescence images of nuclei (DAPI), mature neurons (MAP2) and astrocytes (GFAP) in control and levitated CxSs (a) and StSs (d) at day 6 & 10. Scale bar is 50 *µ*m. Quantification of survival rate of cortical aggregates CxSs (b) and StSs (e) at 3, 6 & 10 days by evaluating the percentage of DAPI-polynucleol+ cells. Each color corresponds to a different experimental batch, with different color map for control and levitated samples. Each circle corresponds to the percentage of healthy cells in one slice, showing dispersion across one sample, while diamonds corresponds to the average percentage for one spheroid. The same graphic chart is conserved further. **CxSs**, DIV3: control (n=4) and levitated (n=3); DIV6: control (n=8) and levitated (n=8); DIV10: control (n=5) and levitated (n=5). **StSS**, DIV3: control (n=5) and levitated (n=4); DIV6: control (n=6) and levitated (n=6); DIV10: control (n=3) and levitated (n=3). Quantification of MAP2+ and GFAP+ cells in control (n=4) and levitated (n=3) CxSs (c) and StSs (f) at 6 & 10 days. Statistical significance assessed with Mann Whitney Wilcoxon test: *p ¡ 0.05, **p ¡ 0.01, ***p ¡ 0.001, ****p ¡ 0.0001.

### 2.3 Impact of acoustic levitation on neural progenitor cells proliferation

Having shown that both CxSs and StSs self-organized in acoustofluidic bioreactors and maintain high viability, their maturation was then evaluated at relevant culture times. First, the immaturity of Neural stem cells (NSCs) was assessed with the specific labeling of Nestin, a marker associated with stemness [34, 35]. The pool of NSCs and neural progenitor cells (NPCs) was defined as directly proportional to the number of Nestin+ cells, with a low percentage corresponding to the exit from cell cycle. In parallel, the proliferation of NPCs was estimated by quantifying Ki67+ cells, a marker for dividing cells [36].

Figure 4.a for cortical spheroids (CxSs) and Figure 4.e for striatal spheroids (StSs) show a nuclear and perinuclear labeling of Nestin as previously observed, with no obvious qualitative difference between control and levitated spheroids for both neuronal subtypes nor between CxSs and StSs. At 3 days of culture, all spheroids displayed a pool of NSCs and NPCs around 20% (Figure 4.b and 4.f). At 6 days of culture, the percentage of Nestin+ cells was more or less stable while it dropped to 13% at 10 days of culture as expected from the maturation of NSCs and NPCs.

**Figure 4:**
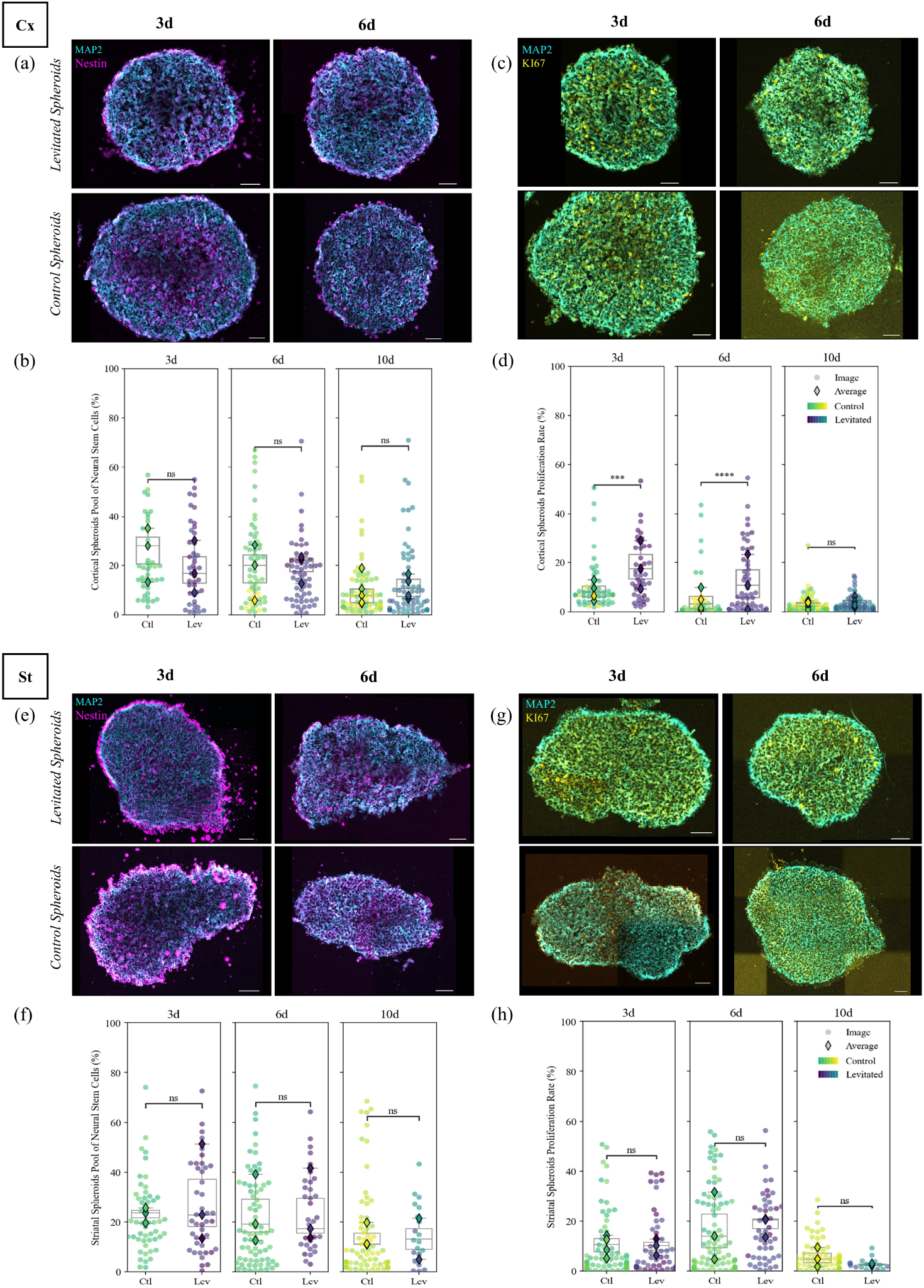
Characterization of cortical (CxSs) and striatal (StSs) spheroids maturation and proliferation at 3 & 6 days. Immunofluorescence images of MAP2 (cyan) and Nestin (magenta) in control and levitated CxSs (a) and StSs (e) at day 3 & 6. Scale bar is 50 *µ*m. Quantification of Nestin+ cells in control (n=3) and levitated (n=3) CxSs (b) and StSs (f). Immunofluorescence images of MAP2 (cyan) and KI67 (yellow) in control and levitated CxSs (c) and StSs (g). Scale bar 50 *µ*m. (d) Quantification of KI67+ cells in control (n=4) and levitated (n=3) CxSs (d) and StSs (h). Statistical significance (Mann Whitney Wilcoxon test) is defined as: *p ¡ 0.05, **p ¡ 0.01, ***p ¡ 0.001, ****p ¡ 0.0001.

We then examined whether the decrease in Nestin+ cells was accompanied by proliferative capacity through Ki67 immunohistochemical analysis (Figure 4.c,d for CxSs and Figure 4.g,h for StSs). In CxSs, the number of proliferating cells was decreasing from 3 to 10 days for both culture conditions. We observed two times more Ki67+ cells in levitated CxSs compared to controls at day 3 (p ¡ 0.001). This effect was kept at day 6 with 3 times more Ki67+ cells in levitated CxSs than in controls (p ¡ 0.0001) while at day 10 almost no KI67+ cells were found. Remarkably, another behavior was found in StSs with a rather low number of KI67+ cells (10%) at 3 days, increasing to 18% at 6 days and decreasing to 5% at 10 days. This shows that acoustic waves transiently and specifically increase the proliferation of cortical NPCs.

### 2.4 Phenotypic Distributions and Synaptic Maturation

The influence of acoustic waves on the differentiation of cortical and striatal cell types was then evaluated using specific markers for phenotypic distribution. Figure 3.c & f show the quantification of neurons and astrocytes for cortical and striatal spheroids respectively. The mean proportion of MAP2+ and GFAP+ cells in CxSs were equivalent between control (48% and 30% resp.) and levitated (38% and 24 % resp.) conditions at 6 days. At later stage, i.e. 10 days of culture, the amount of MAP2+ cells increased to roughly 60% for both conditions, which is consistent with cell cycle exit and maturation of neuronal cell culture. Meanwhile, the number of GFAP+ cells remained quite constant (25% for control and 29% for levitated). Thus, we observed an increase in the ratio of MAP2+/GFAP+, for both control and levitated CxSs from 6 to 10 days.

We observed the same tendency for StSs, with an increase in the number of MAP2+ and GFAP+ cells from 6 to 10 days. At 6 days, the phenotypic distributions were quite similar between the two neuronal subtypes with around 40% of neurons (MAP2+ cells) and 25% of astrocytes (GFAP+ cells). However, at 10 days, the percentage of GFAP+ cells in StSs also increased to roughly 40% in both conditions in opposition to CxSs. Furthermore, we observed a difference in cell phenotypic distributions at 10 days between CxSs and StSs, both in control and levitated conditions, with 10% more MAP2+ cells and almost two times more GFAP+ cells (40%) in StSs compared to CxSs.

We then assessed the excitatory and inhibitory phenotypic index in CxSs and StSs. To elucidate these phenotypes, we verified the status of several markers at the 6-day stage by immunohistochemistry of GAD67 (GABAergic inhibitory neurons) in conjunction with markers of MAP2 (dendrites) and DAPI (nuclear DNA), in levitated and control conditions for cortical (figure 5.a & b) and striatal neurospheres (figure 5.d & e). Quantification of GAD67+ cells showed similar percentage between levitated and control CxSs. Thus, the number of inhibitory neurons in the neuronal population represented by the GAD67+/MAP2+ ratio was equivalent in both culture conditions with 17% in control and 20% in levitated CxSs. For StSs, a vast majority of cells were identified as GAD67+ cells, almost 3 times more than in CxSs. Thus, the percentage of inhibitory neurons over the neuronal population (MAP2+ cells) in StSs reached greater levels (87.2% in control and 89.2% in levitated).

**Figure 5:**
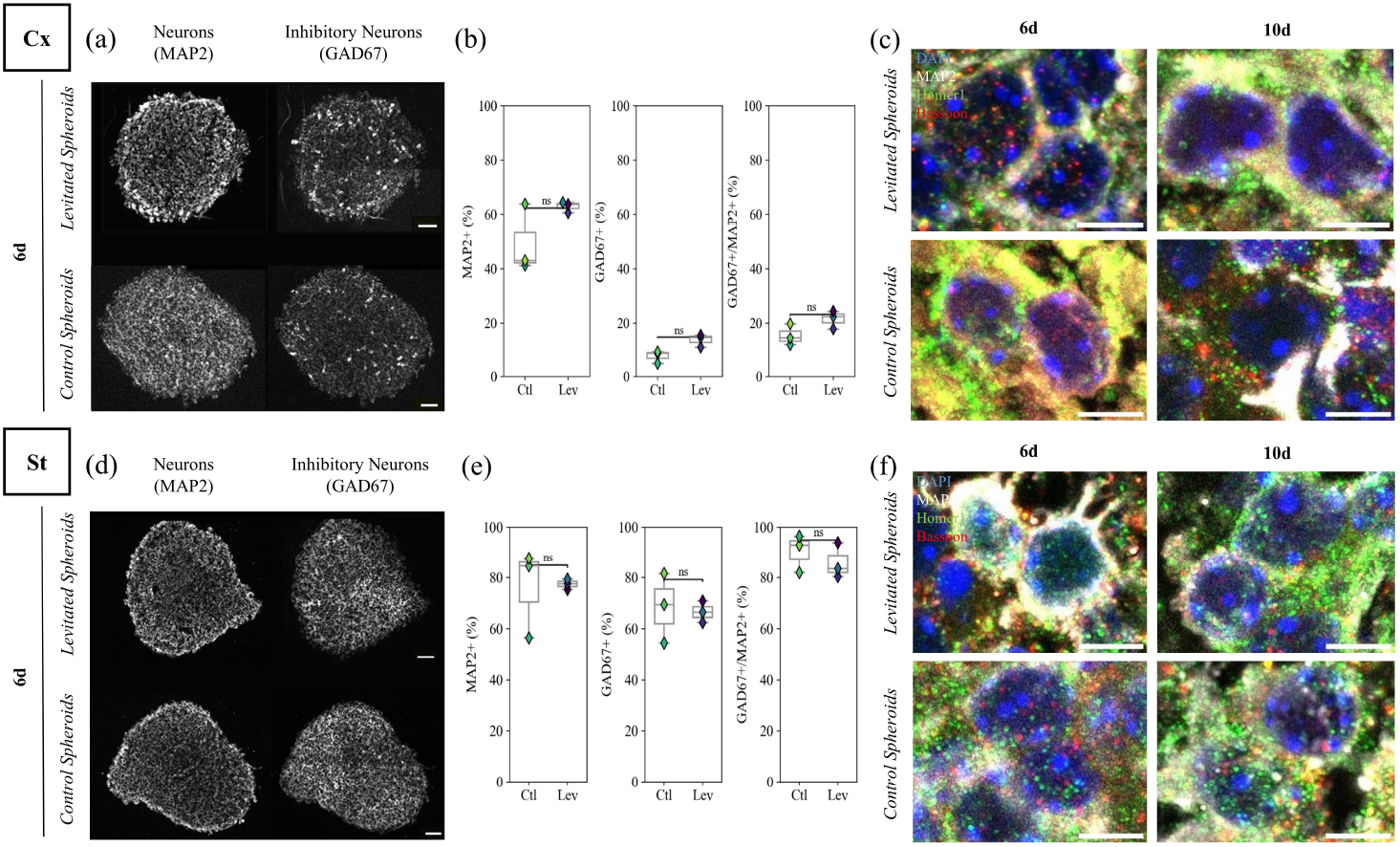
Matured neurospheres showing specific neuronal phenotypes and synaptic connectivity. Characterization of excitatory versus inhibitory neuronal phenotypes of cortical and striatal spheroids, control and levitated, at 6 days. Immunofluorescence images of neurons (MAP2) versus excitatory (VGlut1) and inhibitory (GAD67) neurons showing the phenotypic distributions in spheroids of CxSs (a) & StSs (d) in both culture conditions. Scale bar is 50 *µ*m. Quantification of MAP2+ and GAD67+ cells at day 6 in CxSs (b) & StSs (e); control (n=3) and levitated (n=3). Characterization of pre-& post-synaptic density at 6 & 10 days in cortical (c) and striatal (f) spheroids. Confocal images of immunofluorescence images of MAP2 (white), DAPI (blue), Bassoon (red) and Homer1 (green) in control and levitated StSs at day 6 & 10. Scale bar is 5 *µ*m.

To evaluate synaptic maturation, we performed immunostainings of pre- and post-synaptic markers. Bassoon and Homer1 were chosen as pre- and post-synaptic reporters respectively. In figure 5.c & f, we observe dense pre- and post-synaptic labeling in highly innervated perinuclear areas in both control and levitated cortical and striatal spheroids. This evidence indicates that the spheroids have matured well and formed highly connected neuronal networks.

Altogether, these results demonstrate that acoustic levitation is an amenable technique allowing to self-organize and grow mature and highly connected neuronal spheroids in a scaffold-free environment with a very good viability. After this first validation step, we explored the potential of acoustic levitation to structure and grow complex neuronal spheroids.

## 2.5 Structuration of concentric cortico-striatal assembloids

After showing cortical and striatal neurons cultured in acoustic levitation differentiate and mature well, a single node chip was chosen to build an assembloid made of a striatal core embedded in a cortical shell recapitulating the brain topology (Figure 6.a). This specific chip design was already described in a previous study demonstrating how layers of beads can be arranged concentrically in a cylindrical acoustic cavity [32]. The main application was to study the infiltration of immune cells in glioblastoma tumoroids. Here, to build a Cortico-Striatal Assembloid (CxStAs), both primary neuronal subtypes, cortical and striatal, were extracted and concentrically structured by successive injections of cells, first the core cells (St) and second the shell cells (Cx) - as illustrated in Figure 6.b. However, it remained unclear whether the core-shell structuration obtained in acoustic levitation would be stable over time. To address this question, two structuration protocols were investigated, as illustrated in Figure 7.a. The same tools and methodologies previously described were used for culture and analysis.

**Figure 6:**
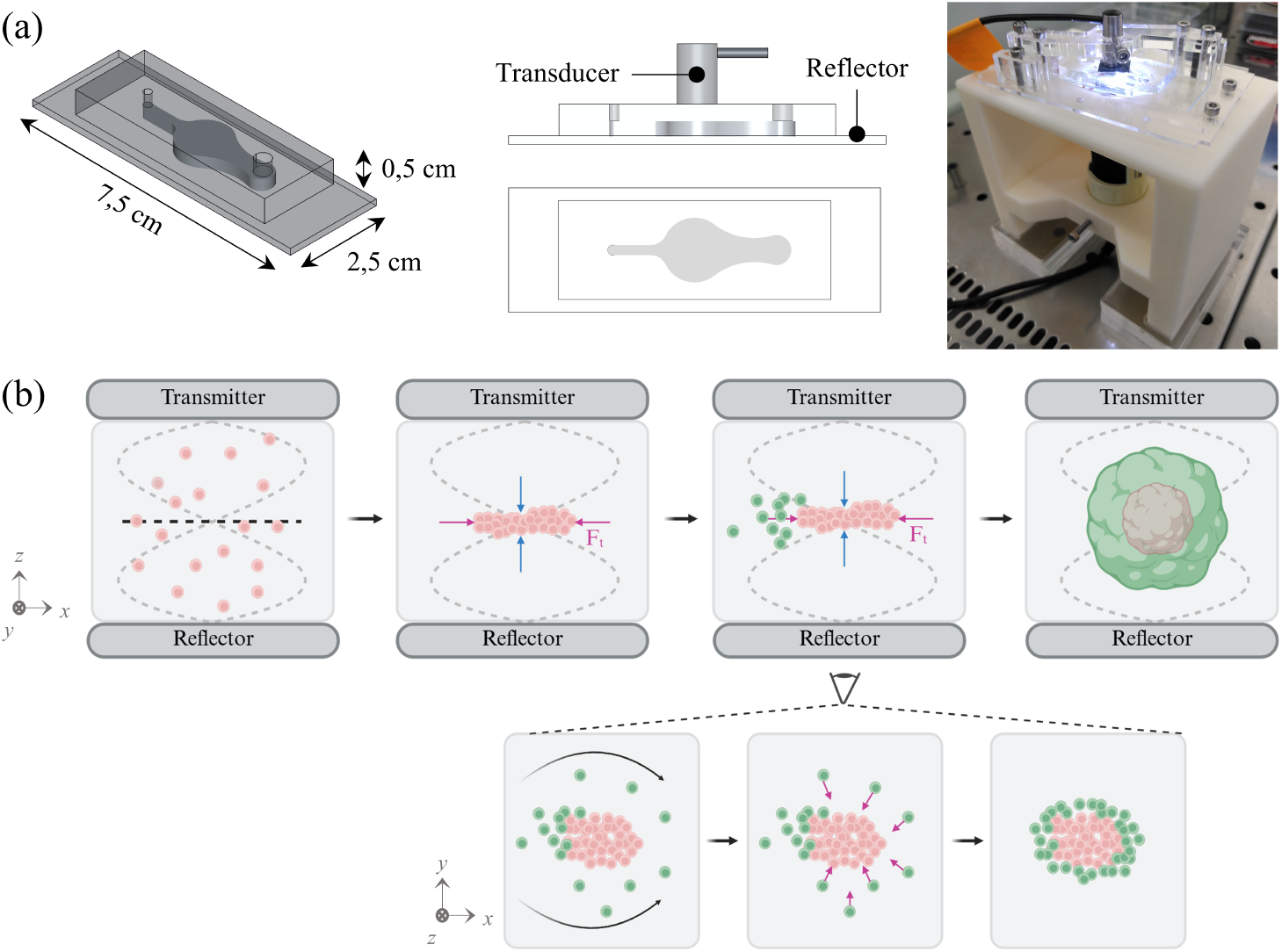
Structuration of a cortico-striatal assembloid. (a) Single-node acoustofluidic chip designed with asymmetrical inlet and outlet for pipette handling and a cylindrical cavity for hydrodynamic forces shaping. The picture on the right shows the 3D printed support holding the camera for axial view imaging inside the hood. (b) Schematics illustrating the formation of a concentrically layered assembloid of two neuronal identities. Striatal neurons (in pink) are first injected and aggregated on the levitation plane inside the single-node acoustofluidic chip. Cortical neurons (in green) are then injected, roughly 10 minutes after striatal neuron injection. Inserts below describe in more details the acoustofluidic phenomenon allowing for the concentric structuration. After injection, cortical neurons first cover the striatal aggregate on the inlet side while some are driven around the aggregate by fluid flows (black arrows), influenced by the horizontal expansion of the cavity. The radial ARF (magenta arrows) then push back the cortical neurons toward the striatal aggregate forming a concentric layer around it. The aggregation takes place for 10 minutes until most cortical cells cover the periphery of the striatal aggregate. Schematics were created with Biorender.com.

**Figure 7:**
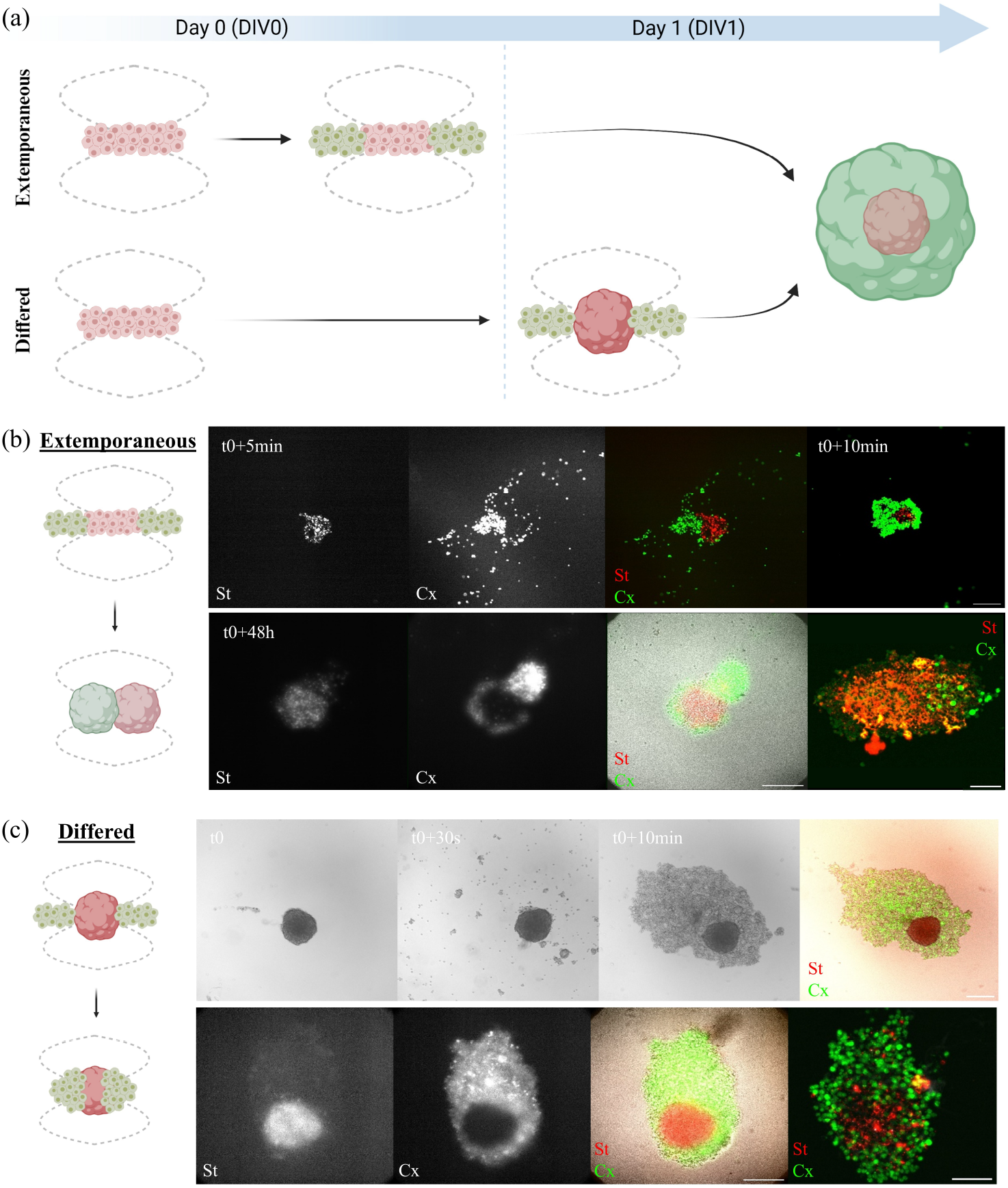
Explored strategies to build Cortico-Striatal Assembloid (CxStAs). (a) Two different strategies were investigated to build concentric topologies. The first protocol consisted in forming the concentric assembloid extemporaneously, by first aggregating striatal cells in acoustic levitation (in red) and immediately surrounding them with cortical cells (in green). The second one was to first form a striatal core spheroid in acoustic levitation and then inject cortical neurons 24 hours later to form an equatorial layer around it. Schematics were created with Biorender.com. (b) For the extemporaneous structuration, both cell types, cortical and striatal neurons, were arranged in the levitation plane in a concentric manner as shown on the small schematic on the left. Pictures of the formation of the CxStAs are presented on the top right with both cell types fluorescently labelled. At the bottom, pictures of the assembloid after 48h in culture are shown. On the right, an immunofluorescence image of a slice of the assembloid highlight the final topology. Scale bars are 100 µm. (c) For the differed structuration, striatal neurons were first aggregated on the levitation plane and allowed to self-organize into a spheroid during 24 hours. At DIV1, cortical neurons were injected in the chip and surrounded the striatal spheroid, forming a concentric annular layer, as shown on the small schematic on the left. Snapshots showing the formation of the CxStAs are presented on the right with both cell types fluorescently labeled. At the bottom, pictures of the assembloid after 48h in acoustic levitation are shown. On the right, an immunofluorescence image of a slice of the assembloid highlight the final topology. Time-lapses of extemporaneous & differed structuration are available on the Supplementary Video 3 & 4 resp. Scale bars are 100 µm.

The first protocol was extemporaneous, meaning that the striatal core layer formed in acoustic levitation was immediately wrapped with a cortical neuronal layer at DIV0, as shown on figure 7.b. At t0+5min, the striatal layer (in red) was already formed at the pressure node with a diameter of roughly 100 µm. At this point, the majority of the striatal cells in suspension near the pressure node had already been attracted towards the aggregate by the radial component of the ARF. Primary cortical neurons (in green) were then injected to surround the striatal core at its periphery. Because of hydrodynamic effects related to the cylindrical geometry of the cavity some of the cells that were passing around the striatal aggregate and moving in the direction of the outlet were pushed back by the transverse component of the ARF towards the striatal core aggregate in levitation at the center of the chamber. Cortical cells coming radially from all directions, they could cover entirely the striatal core layer during the next minutes following the second injection. After 10 min, the structured CxStAs was formed and stable.

The second experimental protocol consisted of forming the striatal core spheroid by acoustic levitation at DIV0 and adding the cortical layer 24 hours later, at DIV1, to improve cell-cell adhesion and territorial segregation (figure 7). The main difference with the first protocole is that the cortical cells are injected after the formation of a striatal spheroid, which should improve the stability of the layering. Indeed, as shown previously in the morphological study, spheroids cultivated in acoustic levitation are cohesive and steady after 24 hours (figure 2). Thus, aggregating the striatal core first and allowing cell adherence could improve the partitioning of the two cell types and prevent rapid migration from one layer to another.

The cortico-striatal assembloids built with the two protocols were then cultured for 2 days in acoustic levitation in a stage incubator allowing their visual monitoring with an inverted microscope. To improve the visualization of the territorial segregation, cortical and striatal cells were labeled with fluorescent dyes which allowed the recording of time-lapse showing the dynamic of the spatial rearrangements of both cell types with live imaging.

As shown in figure 7, the extemporaneous strategy led to the formation of two distinct zones: a striatal spheroid (cells labeled in red) juxtaposed with a cortical spheroid (cells labeled in green). A small diagram on the left represents the final architecture of the CxStAs obtained by this approach. This reorganization was also observed after histoimmunochemical analysis: striatal cells are located in the center, an aggregate of cortical cells touches the striatal nucleus, and a thin layer of cortical cells envelops both spheroids. However, the concentric topology was better preserved with the delayed strategy (figure 7.c). After two days of culture under acoustic levitation, the striatal core spheroid, red-labeled, is surrounded by cortical neurons green-labeled as can be seen both by inverted microscopy and immunohistochemistry images.

Based on this observation, we studied the possibility of transfecting both types of cells, cortical and striatal neurons, with AAVs carrying different fluorophores in order to better elucidate cellular phenotypes and development within the assembloids. A transfection protocol for striatal neurons inside a conventional multiwell plate (DIV0) was developed resulting in the formation of AAV-transfected spheroids after 24 h of culture (DIV1). On the day of structuration (DIV1), cortical neurons were also transfected with AAV-GFP. To build the CxStAs, the mScarlet striatal spheroid was first trapped in acoustic levitation by increasing the voltage of the signal applied to the transducer compared to classical parameters for cell culture (10 *V*_*pp*_). Then, GFP-expressing cortical neurons were injected into the chip and aggregated around the striatal core as previously described.

Figure 8.a shows the CxStAs after 3 days of culture in acoustic levitation (DIV4) with GFP cortical neurons surrounding the mScarlet striatal core spheroid at its equator. The CxStAs was then retrieved and immunostained to highlight viability using DAPI staining. As can be seen from the projection of the full-mount confocal images in figure 8.b, both neuronal subtypes exhibited healthy nuclei and neurites. From this orientation of the assembloid with respect to the confocal microscope objective, GFP-expressing cortical neurons envelop the mScarlet-expressing striatal core more on one side than on the other, although some GFP-Cx can still be seen on the opposite side.

**Figure 8:**
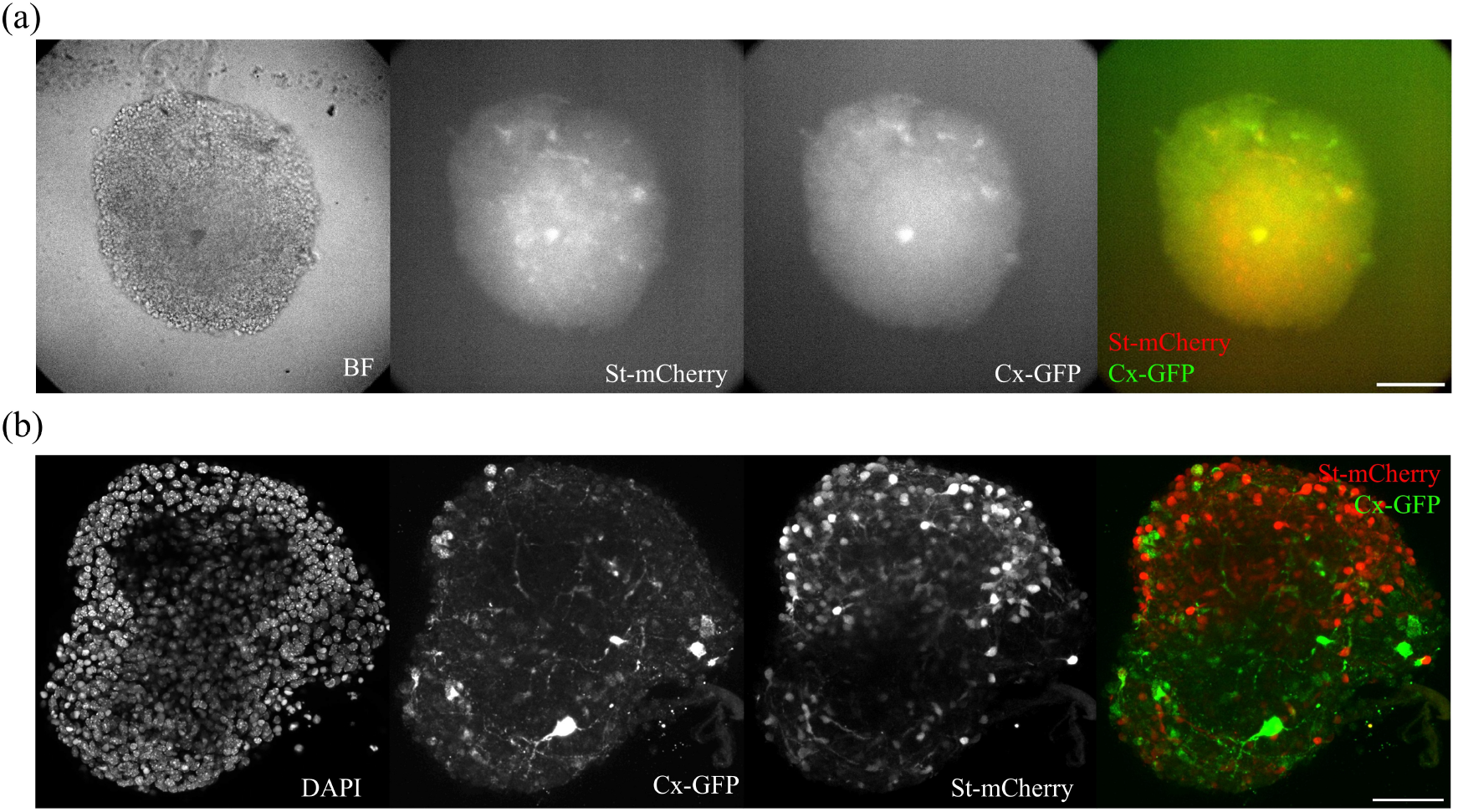
Conserved topology of AAV transfected CxStAs after 3 days of culture. (a) Epifluorescence images of a CxStAs after 3 days of culture in acoustic levitation, composed of cortical cells transfected with AAV-GFP and striatal cells with AAV-mScarlet, structured with the differed strategy. (b) Projections of full-mount confocal images of a CxStAs showing both neuronal subtypes developed neurites, especially Cx-GFP neurons emitting extensions towards the striatal core (mScarlet) at DIV4.

## 3 Discussion

Organ-on-chips and organoids have emerged as powerful models that recapitulate the cellular architecture, biochemical gradients, and physiological responses of human tissues more faithfully than conventional monolayer cultures or animal models [37]. By recapitulating developmental cues and sequences, organoid technologies rely on their capacity to self-assemble into minimalistic organs, accelerating investigations of developmental biology, disease pathogenesis, and drug testing. Yet, their use is limited by the lack of precise control over micro-environmental cues such as shear stress, oxygen tension, and inter-cellular signaling. Advanced technologies such as microfluidic engineering and biosensor interfacing methods allowing control of co-culture are therefore actively explored in order to foster the maturation of 3D cell culture approaches, from rudimentary three-dimensional cell aggregates into sophisticated, physiologically relevant tissue models.

The biofabrication of complex neuronal structures remains a significant challenge using conventional techniques, which often fail to replicate the intricate architecture of native neural networks structures. Our approach leverages acoustic levitation to overcome these limitations by enabling the long-term culture of neurospheres with high viability and by offering the possibility to reconstruct complex neuronal assemblies. Using transverse acoustic force in the acoustic levitation plane, it was possible to successfully engineer concentric assembloids — a novel architecture that mimics the layered organization of neural circuits. This method offers a promising alternative to mechanical positioning or microfluidic approaches, which, while precise, often struggle with spheroid fusion and integration.

Our first observation is that primary neuronal cells rapidly self-organize into 3D neuronal spheroids when cultured in acoustic levitation. This is not specific to neurons, as we previously observed transition from levitated cell monolayers into 3D spheroids with human derived Mesenchymal Stem cells [17] as well as hepatocytes [18] and patient derived gliobastoma [32]. Using the multi-node acoustofluidic chip, we observed differences of shape and surface roughness between cortical and striatal neuronal spheroids. While 3D self-organization in acoustic levitation most probably relies on cell-cell interaction counteracting ARF [18], this may suggest differences in neuron/neuron adhesion forces between striatal and cortical progenitors.

Importantly, we showed that levitated neuronal spheroids exhibit enhanced viability compared to conventional multi-well plate culture methods. This suggests a beneficial effect of acoustic waves on cell survival as previously observed by Rabiet et al. with hepatocytes [19]. Together with the increased survival, we consistently observed a transient increase in the neuronal precursor pool of the levitated spheroids leading to an overall increased proliferation when cultured in acoustic levitation. Our results are also in line with another study that showed that Low Intensity Pulsed UltraSound (LIPUS) were up-regulating the proliferation of 2D cultured Neural Stem Cells (NSCs) derived from iPSCs at a frequency of 1.8 MHz and intensity of 500 *mW*.*cm*^*™*2^ [38]. Indeed, acoustic waves have been shown to influence both differentiation and survival, potentially through modulation of mechanotransduction pathways or global compression forces exerted by ARF [38, 39, 40]. Although we observed an increased proliferation for cortical spheroids, it did not compromise neuronal maturation, resulting in functionally mature neurons comparable to conventionally-grown spheroids. Overall, terminal differentiation was similar between levitated and non-levitated neuronal spheroids, with no modification in the neuron/astrocyte ratio, as well as similar excitatory/inhibitory neuronal populations.

Although this dual effect - increased initial proliferation and adequate terminal differentiation - may suggest accelerated terminal differentiation, it also highlights the potential of acoustic levitation for 3D culture of fragile cell types such as primary rodent neurons. Moreover, thanks to its versatility, the design of our bioreactor paves the way for future refined experiments, for instance, merging multiple spheroids by fine-tuning acoustic parameters [21]. Future work could also explore the impact of pathological pressure levels (e.g., traumatic brain injury models) to further exploit the versatility of this system.

Concentric assembloids represent a significant advance in the reconstruction of neuroanatomical pathways. While previous studies [41] relied on mechanical positioning or on microfluidics for precision, these methods often face challenges in achieving seamless fusions. In contrast, the use of acoustic forces in our bioreactors facilitates the spatial organization of cells, creating a layered architecture that closely resembles native neuronal structures. Although our current approach has imperfections such as ring artifacts, these could be mitigated by more sophisticated systems allowing better spatial control of the acoustic field, for example, through holographic acoustic fields [42]. The work presented here lay the foundations for more complex co-cultures, enabling the study of inter-layer connectivity and functional integration.

Hereafter, the versatility of acoustic levitation-based biofabrication has the potential to engineer increasingly complex co-cultures, combining multiple cell types or incorporating pathological conditions (e.g. TBI-related pressure). The use of multiple sources and more precise acoustic parameters would significantly improve the accuracy of spheroid assembly, thus overcoming current limitations. Ultimately, this platform could become a powerful tool for modeling neuroanatomical pathways, studying pathological mechanisms, and developing therapeutic strategies for neuronal repair.

## 4 Experimental Section

### Microfabrication of Acoustofluidics Chips

First, 3D models of the acoustofluidic chips were designed with Autodesk Inventor. The models were then projected onto a 2D plane with Adobe Illustrator to provide the layout. 2D images were then fed to a laser cutter so that the shape would be cut in a PMMA plate. The thickness of PMMA was chosen to provide adequate height of the chamber for the resonance of ultrasonic wave as described above. PMMA pieces produced were stuck to a plastic container in order to create a mold for acoustic levitation chambers and left to dry 24 hours. Acoustic levitation chambers were made out of PDMS (Sylgard 184, PDMS, Dow Corning) mixed with a curing agent at a ratio of 10:1. The mixture was first degassed in a vacuum chamber and then cast in the dedicated mold. After 2 hours of curing at 70°C, chips were unmolded and fluidic inlet and outlet were punched. Acoustic levitation chamber and microscopic glass slides, serving as reflectors, were treated with O2 plasma cleaner (100% power, 0.6mBar, Diener Electronic). Finally, the acoustofluidic chip was assembled by sticking the glass slide at the bottom of the acoustic levitation chamber. After a short curing time (approximately 2 minutes), distilled water was introduced in the chips to maintain hydrophilicity and chips were sterilized with UV for 30 minutes.

### Primary neurons preparation

SWISS pregnant mice were purchased from Janvier (Le Genest Saint Isle, France). Animal care was conducted in accordance with standard ethical guidelines (U.S. National Institutes of Health publication no. 85–24, revised 1985, and European Committee Guidelines on the Care and Use of Laboratory Animals) and the local, IBPS and UPMC, ethics committee approved the experiments (in agreement with the standard ethical guidelines of the CNRS “Formation ‘a l’Expérimentation Animale” and were approved by the “C2EA-05 Comité d’éthique en experimentation animale Charles Darwin”). Cortices and striata were micro-dissected from 7 to 10 E14 embryos. All steps of dissection were performed in cold Gey’s Balanced Saline Solution (Sigma G9779). Dissected structures were digested with trypsin-EDTA for striata (Gibco) or papaïn for cortices (20U/ML, Sigma 76220) in DMEM. Afer tryspin or papaïn inactivation with FBS, structures were mechanically dissociated with a pipette in presence of DNAse type IV (Sigma D5025). After several rounds of rinsing, cells were re-suspended in culture medium at the desired density. Both neuronal cell types were grown in DMEM glutamax + streptomycin/penicillin (Gibco) + 5% FBS + 2% B27 (Gibco).

### Culture of neurospheres in acoustic levitation

Ultrasonic waves were generated by wide-band ultrasonic transducer (2 MHz TR0213LS, SignalProcessing, Savigny, Switzerland) driven by an arbitrary waveform generator (Handyscope HS5, TiePie Engineering, Sneek, The Netherlands) monitored by a computer. Each output of the signal generator supplied two ultrasonic transducers with a sinusoidal waveform of amplitude 5 Vpp and frequency 2.04 MHz. The acoustic parameters were chosen as a compromise to obtain a good acoustic trapping (choice of the acoustic frequency), while avoiding unwanted phenomena such as acoustic streaming (relatively low voltage). To cultivate primary neurons in acoustic levitation, acoustofluidic chips were first filled with culture medium. The ultrasonic transducer was placed on top of the chip with a thin layer of oil in between the chip and the piezoelectric device to ensure acoustic coupling. Before cells were seeded, the transducer was turned on so that the ultrasonic stationary field would already be stabilized when cells arrived at the center of the chamber to enhance aggregation. Cells were injected at a concentration of 2 million cells/mL in the single-node and 5 million cells/mL in the multi-node acoustofluidic chips, respectively and carefully mixed to homogenize cell distribution inside the levitation chamber. In order to prevent evaporation, the inlet and outlet of the chips were sealed with small PDMS pieces. Finally, acoustic levitation setups containing the chips were placed in an incubator at 37°C, 5% CO2, for several days. Control spheroids were produced by seeding 30 000 cells in ultra-low attachment wells of Nucleon Sphero 96 well-plate.

### Cell-type specific AAV transfection

After dissociation, striatal cells were seeded in 96 well-plates in 100 *µ*L of cell medium containing AAV SD121D (pSyn1-mScarlet, Philippe Ravassard, ICM) but depleted from fetal bovine serum (FBS) and incubated for 4 hours at 37°C. Cells were then rinsed with medium containing FBS three times. They were seeded in the single-node acoustofluidic chip in classic cell culture medium for 24h, allowing them to form an AAV-transfected spheroid. The day of the experiment, cortical cells were incubated for 4 hours at 37°C in cell medium containing AAV GFP 7940 (pSyn1-gfp, Philippe Ravassard, ICM) and without FBS. Cortical cells were then rinsed with medium containing FBS three times before being used.

### Recovery of spheroids

Neuronal spheroids grown under acoustic levitation or in control condition inside low attachment multi well plates were recovered. For levitated spheroids, the transducer was turned off and the chip was removed from the acoustic levitation setup. The spheroid was recovered in 250 µL of medium and fixed in PBS containing 4% paraformaldehyde (PFA, Electron Microscopy Science 15714S) and 4% sucrose (Sigma S9378) for 30 min. To prepare spheroids for OCT snap-freezing, they were put in 15% sucrose for one hour and 30% sucrose for 12 hours minimum. Later, they were embedded in OCT, frozen at −30°C and stored at −80°C. A Cryostat (Leica Biosystems) was then used to cut them in thin slices of 25 µm placed on microscopic glass slides.

### Immunofluorescence staining

Neuronal spheroids sections were washed with PBS for 15 min to remove OCT. Then, they were permeabilized for 5 min with 0.25% Triton (Sigma X100) and 1% bovine serum albumin (BSA, Thermofisher 15561020) in PBS and washed with 10% BSA in PBS for one hour. Primary antibodies were added in PBS with 5% BSA and the samples incubated at 4°C overnight. Samples were then rinsed three times for 5 minutes with PBS and further incubated with the corresponding secondary antibodies in PBS with 5% BSA for 2 hours at room temperature. After three washing with PBS, slides were mounted using Mowiol (475904-M, Sigma) and stored at 4°C. The following primary antibodies were used: DAPI (1050-A, Euromedex, 1/20000), MAP2 (M4403, Sigma, 1/1000), GFAP (PA5-16291, ThermoFisher, 1/1000), B3tubulin (MA1-118, ThermoFisher, 1/1000), Bassoon (ab82958, Abcam, 1/500), Homer1 (160002, Synaptic system, 1/500), GAD67 (MA524909, ThermoFisher, 1/1000). Species-specific secondary antibodies coupled to Alexa 350, 488, 555 or 633 were used (Invitrogen, 1/1000) to visualize bound primary antibodies.

### Image acquisition

Images were acquired with an epifluorescence Axio-observer Z1 (Zeiss) equipped with a cooled CCD camera (CoolsnapHQ2, Roper Scientifc) or a confocal microscope (Zeiss LSM980 Fast-Airyscan 2). The epifluoresence microscope was controlled with Metamorph software (Molecular Imaging) and images were analyzed using ImageJ software (ImageJ, U.S. National Institutes of Health, Bethesda, Maryland, USA).

### Image analysis

#### Definition of Morphodescriptors

To quantify the time evolution of the shape and size of spheroids during cultivation in acoustic levitation, a dedicated algorithm was coded on *Matlab* software (R2022a). The first step consists in binarizing the image to obtain the contour (figure 9.a) then the area *A*_*s*_(*t*) of the spheroid at time *t* (figure 9.b). As the picture is a side view of the spheroids, this area corresponds to the projection of the aggregate onto the vertical plane. Several geometrical parameters can then be determined from *A*_*s*_. First, the equivalent diameter *D*_*e*_ is computed using eq. 1:

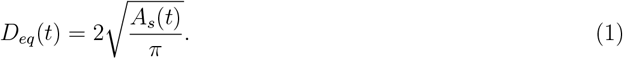

**Figure 9:**
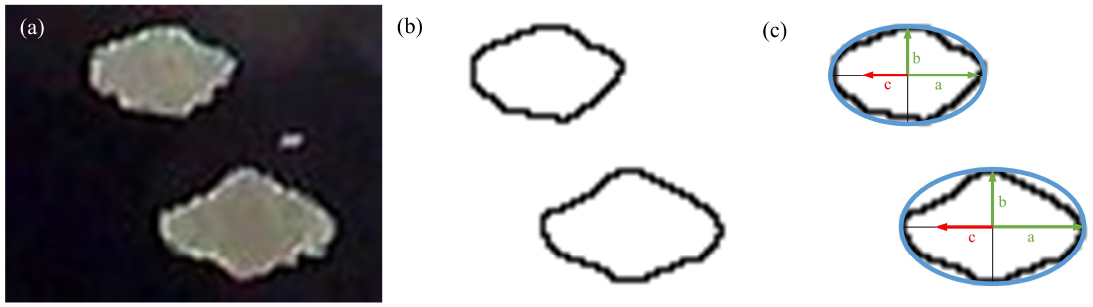
Description of Morphodescriptors. From snapshots of aggregates taken from the side (a), contours are extracted through image processing (b). Based on these contours, 3 parameters could be computed to describe the shape of spheroids: Equivalent diameter, circularity and eccentricity (c).

It gives an estimate of the mean diameter of an equivalent circle, the aggregate being far from circular, especially during the first hours.

Another interesting parameter is the instantaneous circularity *Circ*(*t*) of the aggregate (figure 9.c). It is calculated using eq. 2:

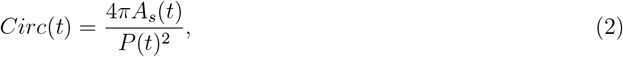

where *P* (*t*) is the perimeter of the measured area *A*_*s*_(*t*). One can see that a perfect circle will have a circularity equal to 1, while the circularity of a flattened ellipse will tend towards 0.

The last parameter is the eccentricity *Ecc*(*t*) of the aggregate, if it is considered as an ellipse, defined by the eq. 3:

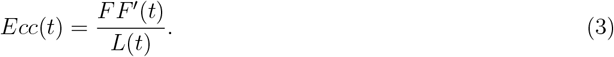

If the aggregate is considered as an ellipse then it is possible to determine *FF*^*′*^(*t*), which corresponds to the distance between the two foci of the ellipse and *L*(*t*) which corresponds to the length of its principal axis. The value of eccentricity ranges from 0 for a perfect circle to 1 for a segment. The instantaneous eccentricity of the aggregates will then vary over time from close to 1 (flat aggregate) to approximately 0 (sphere).

#### Analysis of immunofluorescence images

Custom made routines based on results obtained by manual classification were developed in FIJI for each immunostaining marker. The segmentation of nuclei and the extraction of cell positions were based on fluorescent nucleic acid labeling (DAPI) images of spheroids slices. A Plugin developed by Schmidt and Weigert [43, 44] called StarDist was used for segmentation. Based on previous works [45, 46, 47], we defined poly-nucleolated nuclei as alive and mono-nucleolated, condensed, nuclei with high intensity as dead. Experimenters with relevant experience were asked to manually sort more than 20 segmented nuclei as either alive or dead in 5 different fields of view. Criteria based on the size, mean and max intensity of nuclei as well as number and size of nucleoli were extracted from this manual classification and then used to establish the automatic classification. For the differentiation, proliferation and phenotypic distribution analysis, the classification of positively stained cells was made only on Region Of Interests (ROI) previously identified as alive cells by DAPI analysis. Different thresholds to define ROI as positively stained cells were evaluated manually for each marker. To decrease experimental noise, the mean and max intensity of each ROI were normalized by the mean intensity of the whole image in order to bring out a valid threshold defining staining i.e. positively stained cells. For nuclear and perinuclear labeling (Nestin and KI67), thresholding was applied on the same ROIs defined by DAPI segmentation. For somatic labeling (MAP2, GFAP, GAD67), ROIs were modified to form a ring shape around the nuclei. Mann-Whitney-Wilcoxon two-sided tests were performed on the different images in between conditions to extract p values.

## Supporting Information

Supporting Information is available from the Wiley Online Library or from the author.

## Acknowledgements

This work was supported by the Ecoles Doctorales Sciences mécaniques, acoustique, électronique et robotique de Paris - ED 391 and Cerveau, Cognition, Comportement - ED 158 of Sorbonne University and the ANR Octobol (ANR-23-CE52-0010).

## Ethics Statement

Animal care was conducted in accordance with standard ethical guidelines (U.S. National Institutes of Health publication no. 85–24, revised 1985, and European Committee Guidelines on the Care and Use of Laboratory Animals) and the local, IBPS and UPMC, ethics committee approved the experiments (in agreement with the standard ethical guidelines of the CNRS “Formation à l’Expérimentation Animale” and were approved by the “C2EA-05 Comité d’éthique en experimentation animale Charles Darwin”).

## Data Availability Statement

All data, code, and materials that support the findings of this study are available in [Repository Name] at [DOI or persistent link]. For any additional information or requests, please contact the corresponding author at.

## References

[1] J. Kim, B.-K. Koo, J. A. Knoblich, Nature Reviews Molecular Cell Biology 2020, 21, 10 571, number: 10 Publisher: Nature Publishing Group.

[2] E. Karzbrun, A. Kshirsagar, S. R. Cohen, J. H. Hanna, O. Reiner, Nature Physics 2018, 14, 5 515.

[3] R. Habibey, J. E. Rojo Arias, J. Striebel, V. Busskamp 122, 18 14842, publisher: American Chemical Society.

[4] Y.-C. E. Li, Y. A. Jodat, R. Samanipour, G. Zorzi, K. Zhu, M. Hirano, K. Chang, A. Arnaout, S. Hassan, N. Matharu, A. Khademhosseini, M. Hoorfar, S. R. Shin, Biofabrication 2020, 13, 1 10.1088/1758.

[5] J. Kajtez, M. F. Wesseler, M. Birtele, F. R. Khorasgani, D. Rylander Ottosson, A. Heiskanen, T. Kamperman, J. Leijten, A. Martínez-Serrano, N. B. Larsen, T. E. Angelini, M. Parmar, J. U. Lind, J. Emnéus, Advanced Science 2022, 2201392.

[6] J. Andersen, O. Revah, Y. Miura, N. Thom, N. D. Amin, K. W. Kelley, M. Singh, X. Chen, M. V. Thete, E. M. Walczak, H. Vogel, H. C. Fan, S. P. Paşca, Cell 2020, 183, 7 1913.

[7] J.-i. Kim, Y. Miura, M.-Y. Li, O. Revah, S. Selvaraj, F. Birey, X. Meng, M. V. Thete, S. D. Pavlov, J. Andersen, A. M. Paşca, M. H. Porteus, J. R. Huguenard, S. P. Paşca, Human assembloids reveal the consequences of CACNA1G gene variants in the thalamocortical pathway, 2023, Pages: 2023.03.15.530726 Section: New Results.

[8] J.-i. Kim, K. Imaizumi, O. Jurjut,, K. W. Kelley, D. Wang, M. V. Thete, Z. Hudacova, N. D. Amin, R. J. Levy, G. Scherrer, S. P. Pas, ca, Nature 2025, 1–11, publisher: Nature Publishing Group.

[9] H. Bruus, Theoretical microfluidics, Number 18 in Oxford master series in physics. Oxford University Press, OCLC: ocn156817008.

[10] M. Settnes, H. Bruus, Physical Review E 2012, 85, 1 016327, publisher: American Physical Society.

[11] E. Christakou, M. Ohlin, B. Önfelt, M. Wiklund, Lab on a Chip 2015, 15, 15 3222.

[12] K. Olofsson, V. Carannante, M. Ohlin, T. Frisk, K. Kushiro, M. Takai, A. Lundqvist, B. Önfelt, M. Wiklund, Lab on a Chip 2018, 18, 16 2466.

[13] B. Kang, J. Shin, H.-J. Park, C. Rhyou, D. Kang, S.-J. Lee, Y.-s. Yoon, S.-W. Cho, H. Lee, Nature communications 2018, 9, 1 5402.

[14] C. Bouyer, P. Chen, S. Güven, T.T. Demirtaş, T. Nieland, F. Padilla, U. Demirci, Advanced materials (Deerfield Beach, Fla.) 2015, 28, 1 161.

[15] D. V. Deshmukh, P. Reichert, J. Zvick, C. Labouesse, V. Künzli, O. Dudaryeva, O. Bar-Nur, M. W. Tibbitt, J. Dual, Advanced Functional Materials 2022, 32, 30 2113038.

[16] T. Miao, K. Chen, X. Wei, B. Huang, Y. Qian, L. Wang, M. Xu, International Journal of Bioprinting 2023, 9, 4 733.

[17] N. Jeger-Madiot, L. Arakelian, N. Setterblad, P. Bruneval, M. Hoyos, J. Larghero, J.-L. Aider, Scientific Reports 2021, 11, 1 8355.

[18] L. Rabiet, L. Arakelian, N. Jeger-Madiot, D. García, J. Larghero, J.-L. Aider, Biotechnology and Bioengineering 2024, 121.

[19] L. Rabiet, N. Jeger-Madiot, D. García, L. Tosca, G. Tachdjian, S. Kellouche, R. Agniel, J. Larghero, J.-L. Aider, L. Arakelian, Scientific Reports 2024, 14.

[20] S. Li, P. Glynne-Jones, O. G. Andriotis, K. Y. Ching, U. S. Jonnalagadda, R. O. Oreffo, M. Hill, R. S. Tare, Lab on a Chip 2014, 14, 23 4475.

[21] N. Jeger-Madiot, X. Mousset, C. Dupuis, L. Rabiet, M. Hoyos, J.-M. Peyrin, J.-L. Aider, The Journal of the Acoustical Society of America 2022, 151, 6 4165.

[22] Z. Ao, H. Cai, Z. Wu, J. Ott, H. Wang, K. Mackie, F. Guo, Lab on a chip 2021, 21, 4 688.

[23] K. Olofsson, V. Carannante, M. Takai, B. Önfelt, M. Wiklund 12, 3 329, number: 3 Publisher: Multidisciplinary Digital Publishing Institute.

[24] G. Whitworth, W. T. Coakle 91, 1 79, place: Woodbury, NY Publisher: Acoustical Society of America.

[25] G. Dumy, N. Jeger-Madiot, X. Benoit-Gonin, T. E. Mallouk, M. Hoyos, J.-L. Aider, Microgravity Science and Technology 2020, 32, 6 1159.

[26] V. Bjerknes, Fields of force: supplementary lectures, applications to meteorology; a course of lectures in mathematical physics delivered December 1 to 23, 1905, 1. Columbia University Press, 1906.

[27] Doinikov, Journal of Fluid Mechanics 2001, 444 1.

[28] Garcia-Sabaté A. Castro, M. Hoyos, R. González-Cinca, The Journal of the Acoustical Society of America 2014, 135, 3 1056.

[29] G. T. Silva, H. Bruus, Physical Review E 2014, 90, 6 063007.

[30] R. Barnkob, I. Iranmanesh, M. Wiklund, H. Bruus, Lab on a Chip 2012, 12, 13 2337.

[31] L. Bellebon, H. R. Sugier, J. Larghero, J. Peltzer, C. Martinaud, M. Hoyos, J.-L. Aider, Frontiers in Physics 2022, 10.

[32] M. Xavier, K. Shuake, D. Chloé, J.-M. Nathan, D. Virgile, I. Ahmed, E.-H. A. Elias, C. Hervé, J. Marie-Pierre, A. Jean-Luc, et al., bioRxiv 2025, 2025–02.

[33] Y. A. Hannun, R.-M. Boustany, Apoptosis in Neurobiology, CRC Press, 1998, google-Books-ID: pabeTqlXO 0C.

[34] U. Lendahl, L. B. Zimmerman, R. D. McKay, Cell 1990, 60, 4 585.

[35] Wiese, A. Rolletschek, G. Kania, P. Blyszczuk, K. V. Tarasov, Y. Tarasova, R. P. Wersto, K. R. Boheler, A. M. Wobus, Cellular and molecular life sciences: CMLS 2004, 61, 19–20 2510.

[36] N. Kee, S. Sivalingam, R. Boonstra, J. M. Wojtowicz, Journal of Neuroscience Methods 2002, 115, 1 97.

[37] N. J. Fedorchak, N. Iyer, R. S. Ashton 111 52.

[38] Y. Lv, P. Zhao, G. Chen, Y. Sha, L. Yang 35, 12 2201.

[39] J. Wen, X. Deng, C. Huang, Z. An, M. Liu 12, 11 1931.

[40] T. Papp, Z. Ferenczi, B. Szilagyi, M. Petro, A. Varga, E. Kókai, E. Berenyi, G. Olah, G. Halmos, P. Szucs, Z. Meszar 16.

[41] Y. Miura, M.-Y. Li, O. Revah, S.-J. Yoon, G. Narazaki, S. P. Pas, ca, Nature Protocols 2022, 17, 1 15, number: 1 Publisher: Nature Publishing Group.

[42] Marzo, S. A. Seah, B. W. Drinkwater, D. R. Sahoo, B. Long, S. Subramanian, Nature communications 2015, 6, 1 8661.

[43] U. Schmidt, M. Weigert, C. Broaddus, G. Myers, In A. F. Frangi, J. A. Schnabel, C. Davatzikos, Alberola-López, G. Fichtinger, editors, Medical Image Computing and Computer Assisted Intervention – MICCAI 2018, Lecture Notes in Computer Science. Springer International Publishing, Cham, ISBN 978-3-030-00934-2, 2018 265–273.

[44] M. Weigert, U. Schmidt, R. Haase, K. Sugawara, G. Myers, In Proceedings of the IEEE/CVF winter conference on applications of computer vision. 2020 3666–3673.

[45] Shiels, N. M. Adams, S. A. Islam, D. A. Stephens, P. S. Freemont 3, 7 e138, publisher: Public Library of Science.

[46] Ferro, T. Mestre, P. Carneiro, I. Sahumbaiev, R. Seruca, J. M. Sanches, Laboratory Investigation 2017, 97, 5 615, number: 5 Publisher: Nature Publishing Group.

[47] P. Rana, A. Sowmya, E. Meijering, Y. Song 11, 1 3364, number: 1 Publisher: Nature Publishing Group.

